# A Gut Microbial Peptide and Molecular Mimicry in the Pathogenesis of Type 1 Diabetes

**DOI:** 10.1101/2020.10.22.350801

**Authors:** Khyati Girdhar, Qian Huang, I-Ting Chow, Claudia Brady, Amol Raisingani, Patrick Autissier, Mark A. Atkinson, William W. Kwok, C. Ronald Kahn, Emrah Altindis

## Abstract

Type 1 Diabetes (T1D) is an autoimmune disease characterized by the destruction of pancreatic β-cells. One of the earliest aspects of this process is development of autoantibodies and T-cells directed at an epitope in the B-chain of insulin (insB:9-23). Analysis of microbial protein sequences with homology to insB:9-23 sequence revealed 17 peptides showing >50% identity to insB:9-23. Of these, one peptide, found in the normal human gut commensal *Parabacteroides distasonis*, activated both human T cell clones from T1D patients and T-cell hybridomas from non-obese diabetic (NOD) mice specific to insB:9-23. Immunization of NOD mice with *P. distasonis* insB:9-23 peptide mimic or insB:9-23 peptide verified immune cross-reactivity. Colonization of female NOD mice with *P. distasonis* accelerated the development of T1D, increasing macrophages, dendritic cells and destructive CD8+ T-cells, while decreasing FoxP3+ regulatory T-cells. Western blot analysis identified *P. distasonis* reacting antibodies in sera of NOD mice colonized with *P. distasonis* and human T1D patients. Furthermore, adoptive transfer of splenocytes from *P. distasonis* treated mice to NOD/SCID mice enhanced disease phenotype in the recipients. Finally, analysis of human infant gut microbiome data revealed that exposure of infants to *P. distasonis* may modulate disease pathogenesis. Taken together, these data demonstrate the potential role for an insB:9-23-mimimetic peptide from gut microbiota as a molecular trigger or modifier of T1D pathogenesis.

**SIGNIFICANCE STATEMENT:** In Type 1 diabetes (T1D), immune cells destroy pancreatic β-cells. The trigger of this response, however, is unknown. Some sequences (epitopes) in the insulin molecule form a major target for this autoimmune response. We have identified a sequence in a human gut bacterium that can mimetic this insulin epitope. Immune cells specific to insulin cross-react with this bacterial mimetic. Further, this bacterium can accelerate diabetes onset in a mouse model of T1D, inducing destructive and decreasing protective immune cells. We found this mimetic in the gut of children developing T1D. Furthermore, T1D patients have a stronger immune response to this bacterium compared to healthy individuals. Taken together, this bacterial mimetic in human gut has the potential to trigger/modify T1D onset.

## 1. INTRODUCTION

Type 1 diabetes (T1D) is an autoimmune disease characterized by selective destruction of pancreatic β-cells by autoreactive T-cells(1). Genome-wide association studies (GWAS) have identified 152 genetic regions that influence the risk of developing T1D(2); however, multiple studies have shown that the incidence rate of T1D in children is rising at rates exceeding what can be explained on a genetic basis alone(3). Indeed, even among identical twins, the concordance of T1D is only 65%(4). Likewise, there is a six-fold difference in incidence of T1D in neighboring regions of Karelia in Russia and Finland, despite the very similar genetic background of the inhabitants(5).

Various environmental factors have been studied as potential modifiers or triggers of the autoimmune response in T1D including diet, birth mode, infection, and antibiotics. Viral infections have been suggested to play a role in T1D pathogenesis, but most of these viruses have been proposed to act by direct infection of the β-cell(6). Recently, attention has been focused on the gut microbiome as a potential disease modifier through its effects on metabolite composition, intestinal permeability, and regulation of the immune response in subjects with T1D(7, 8). However, the exact environmental modifiers and how they might affect T1D pathogenesis remain largely unknown(9).

One of the earliest markers of T1D is the development of islet autoantibodies(10). These autoantibodies target several autoantigens including insulin, glutamic acid decarboxylase (GAD), insulinoma-associated autoantigen-2 (IA-2), zinc transporter-8 (ZnT8), and an islet-specific glucose-6-phosphatase catalytic subunit related protein (IGRP or G6PC2)(11). Among these, insulin autoantibodies (IAA) are usually the first to be detected, and insulin is the only autoantigen exclusively expressed by β-cells(12). In humans, IAA may develop years before the onset of overt diabetes(13) and are especially prominent in early-onset T1D(14). They also show a significant correlation with the rate of progression from prediabetes to overt disease(15). More importantly, insulin or insulin-derived peptides are a target of disease pathogenetic T-cells in both humans with T1D and in the most established murine model, the NOD mouse. In NOD mice, over 90% of the anti-insulin T-cell clones target a single 15-amino acid peptide corresponding to the insulin B-chain 9-23 sequence (insB:9-23)(15). InsB:9-23 specific T-cells have also been identified in islets(16) and peripheral blood lymphocytes of T1D patients(17, 18).

In the present study, we hypothesized that exposure to a microbial peptide that resembles the insulin epitope, insB:9-23, could stimulate or modify the autoimmune response initiating T1D(19). To address this hypothesis, we analyzed bacterial, viral and fungal genome databases to identify microbial proteins that have >50% sequence homology to human insB:9-23. Of these, 17 were synthesized and tested for ability to activate insB:9-23 specific T-cells. Herein, we demonstrate that one of these peptides, a peptide from the gut commensal organism *Parabacteroides distasonis*, could activate both human and NOD mouse insB:9-23-specific T-cells *ex-vivo* to the same extent as the human insulin B:9-23 peptide. This bacterial peptide also cross-reacts with the immune cells obtained from the mice immunized with the human insB:9-23 peptide. Furthermore, administration of *P. distasonis* bacteria by oral gavage accelerated T1D progression in NOD mice *in vivo* by stimulating innate immune cells and CD8+ T-cells and decreasing regulatory T-cells. Analysis of human gut microbiota datasets from the longitudinal DIABIMMUNE study revealed the presence of this bacterial peptide in children who later developed autoantibodies to insulin itself. Taken together, these studies demonstrate that an insulin B:9-23-like epitope in normal gut microbiota may mimic the native insulin peptide and has potential to play a role in onset or progression of T1D.

## 2. RESULTS

### 2.1 A *P. distasonis* insB:9-23-like peptide stimulates NOD mouse hybridomas and human T-cell clones specific to insB:9-23

To determine if any microbe might have DNA encoding proteins with sequences resembling the dominant T-cell epitope involved in the autoimmune response to the insulin B:9-23 peptide linked to T1D (SHLVEALYLVCGERG)(15–18), we used BlastP to search NCBI databases for predicted proteomes of all sequenced viruses (taxid:10239), bacteria (taxid:2), and fungi (taxid:4751). We identified 47 microbial peptides with over eight residues identical to the insulin peptide **(Supplementary Table S1**). Among these predicted microbial peptides, we selected 17 bacterial and viral peptides that contained the largest number of previously identified residues in the insB:9-23 peptide critical for this interaction(18) to be synthesized (**Figure 1A**).

**Figure 1.**
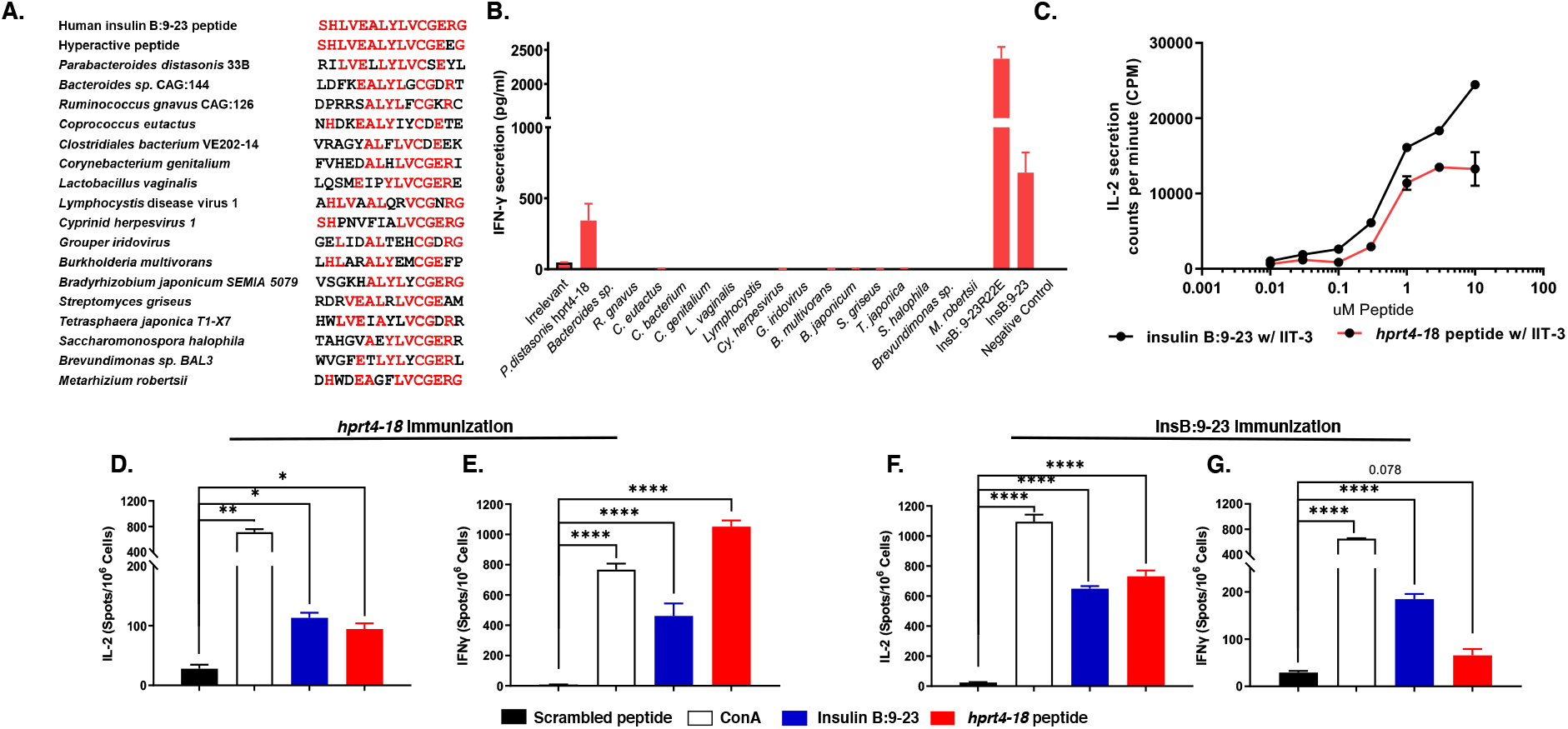
*P. distasonis hprt4-18* peptide mimic can stimulate both human and NOD mice T cells specific to insB:9-23. **(A)** Sequence alignment of 17 microbial peptides tested in this study. The amino acids highlighted in red are identical to insB:9-23. (**B)** Treatment of insB:11-23-specific human T-cell clones with each of selected 17 microbial peptides, where, negative control and irrelevant peptides were used as the negative controls, ins B9-23 and insB:9-23 R^22E^ were used as positive controls. ELISpot was used to determine IFN-γ secretion *(P<0.05)* (n=3). **(C)** Dose response of IIT-3 hybridomas to insB:9-23 (black) or *hpt4-18* peptides to (red) where peptides were presented to hybridomas as covalently linked to I-A^g7^ expressed on macrophages (C3g7 cells). Data were presented as IL-2 induced proliferation in CTLL-2 cells. **(D-G)** Mice (n=3/group) were immunized with either *hprt4-18* peptide (**D-E**) or insB:9-23 peptide (**F-G**) after 7 days of immunization, popliteal lymph nodes were stimulated either by *hprt4-18* peptide, insB:9-23, ConA (positive control) or scrambled peptide (negative control). Secretion of IL-2 (**E & G**) and IFN-γ **(D & F)** were determined by ELISpot. All samples in each panel are biologically independent. Data were expressed as means ± SEM. *\*P<0.05*, *\*\*P<0.01*, *\*\*\*\*P<0.0001*. Statistical analysis was performed by one-way ANOVA using Dunnett’s multiple comparison test.

These 17 microbial insB:9-23-like peptides, as well as a negative control (irrelevant/scramble) and a positive control (insB:9-23), were then tested for their ability to stimulate an insB:11-23-specific human T-cell clone isolated from a T1D patient(18) (**Figure 1B**). In addition, we included a second positive control, a variant of the insB:9-23 peptide in which R at position 22 is substituted by E (insB:9-23R^22E^) and which has been shown to be even more potent than native insB:9-23 in stimulating insB:9-23 hybridomas from NOD mice(20) and human T-cell clones(18). As expected, in the human T-cell assay, both positive controls stimulated interferon-gamma (IFN-γ) secretion with insB:9-23R^22E^ being more active than the wild type sequence. Among the 17 microbial peptides tested, only one could stimulate human T-cells to produce the IFN-γ (**Figure 1B**). This peptide (*hprt4-18*) represents amino acids 4-18 in the N-terminus of the hypoxanthine phosphoribosyltransferase (*hprt*) protein of *Parabacteriodes distasonis* 33B and D13 strains (formerly known as *Bacteroides sp. 2_1_*33B*/ Parabacteroides sp.* D13)(21). Interestingly, *P. distasonis* 33B and D13 are the only organisms in the NCBI dataset (as of October, 2021) that possess this insB:9-23 mimic sequence in their genomes. Next, we tested the same set of microbial insB:9-23-like peptides and controls on NOD IIT-3 T-cell hybridomas, previously shown to recognize the insB:9-23 epitope(22). In this assay, C3g7 cells that have high expression of MHC-II I-A^g7^ were used as antigen-presenting cells (APCs) and treated with each peptide at concentrations from 0.01 to 10μM. Co-culturing of these cells with IIT-3 T-cell hybridomas stimulated interleukin-2 (IL-2) secretion only in presence of insB:9-23 and the *hprt4-18* peptide, and produced similar dose-response curves, with the bacterial peptide being only slightly less potent **(Figure 1C**). Thus, both human T-cell clones and NOD mouse T-cell hybridomas specific to insB:9-23 were reactive to *hprt4-18*, raising the possibility that *hprt4-18* may have the potential to modulate the development of T1D.

### 2.2 *P. distasonis hprt4-18* peptide stimulates a T-cell response to insB:9-23 in vivo

To further explore the cross-reactivity of the microbial and insulin-derived peptide sequences, NOD mice were immunized with either insB:9-23 peptide or the *hprt4-18* peptide in Complete Freund’s Adjuvant (CFA). After 7 days, lymphocytes were isolated from the popliteal lymph nodes and stimulated with either insB:9-23, *hprt4-18* or control peptides. T-cell activation was assessed using an enzyme-linked immunospot (ELISpot) assay. Consistent with the in-vitro results above, lymphocytes from mice immunized with the *hprt4-18* peptide exhibited a strong immune response to both *hprt4-18* peptide and insB:9-23 as measured by IFNγ and IL-2 secretion (**Figure 1D & 1E**). Conversely, lymphocytes from mice immunized with insB:9-23 peptide showed a strong response to both insB:9-23 peptide and the microbial *hprt4-18* peptide (**Figure 1F & 1G**). Taken together with the assays using the hybridoma T-cell lines, these data indicate that the microbial peptide *hprt4-18* strongly cross-react with the recognition and signaling machinery for human insB:9-23 in stimulating an immune response.

### 2.2 *P. distasonis* colonization accelerates diabetes onset in NOD mice

Based on this cross-reactivity, we hypothesized that *P. distasonis* colonization and potential exposure to the microbial *hprt4-18* peptide could trigger and/or modify the immune response and stimulate autoimmunity in NOD mice. To test this hypothesis, both male and female NOD mice were orally gavaged with a saline suspension of *P. distasonis* (10**^8^** CFU/mouse/day) or saline itself daily, starting at 4 weeks of age (i.e., after weaning) and followed for 30 weeks (**Figure 2A**, n=40/group). Two weeks after the last oral gavage, qPCR using fecal DNA samples revealed significant levels of *P. distasonis* in the feces of the treated mice, whereas neither male nor female control NOD mice housed under specific pathogen-free (SPF) conditions had *P. distasonis* in their gut microbiome (**Figure 2B**). At 12 weeks of age, when mice are in the pre-diabetic stage, five mice from each group were randomly selected and examined histologically for insulitis. In the *P. distasonis* colonized female NOD mice there was more than 2-fold increase in severe insulitis scores as compared to controls (**representative images in 2C, Figure 2D**). Consistent with the increase in insulitis, female NOD mice subjected to *P. distasonis* colonization showed significantly accelerated onset of T1D with only 19.5% disease-free in the *P. distasonis* colonized group versus 42% disease-free in the control group (**Figure 2E**). This effect was specific to the female mice, with no differences in insulitis scores or T1D incidence in male NOD mice with *P. distasonis* colonization (**Supplementary Figure S1A-B)**.

**Figure 2.**
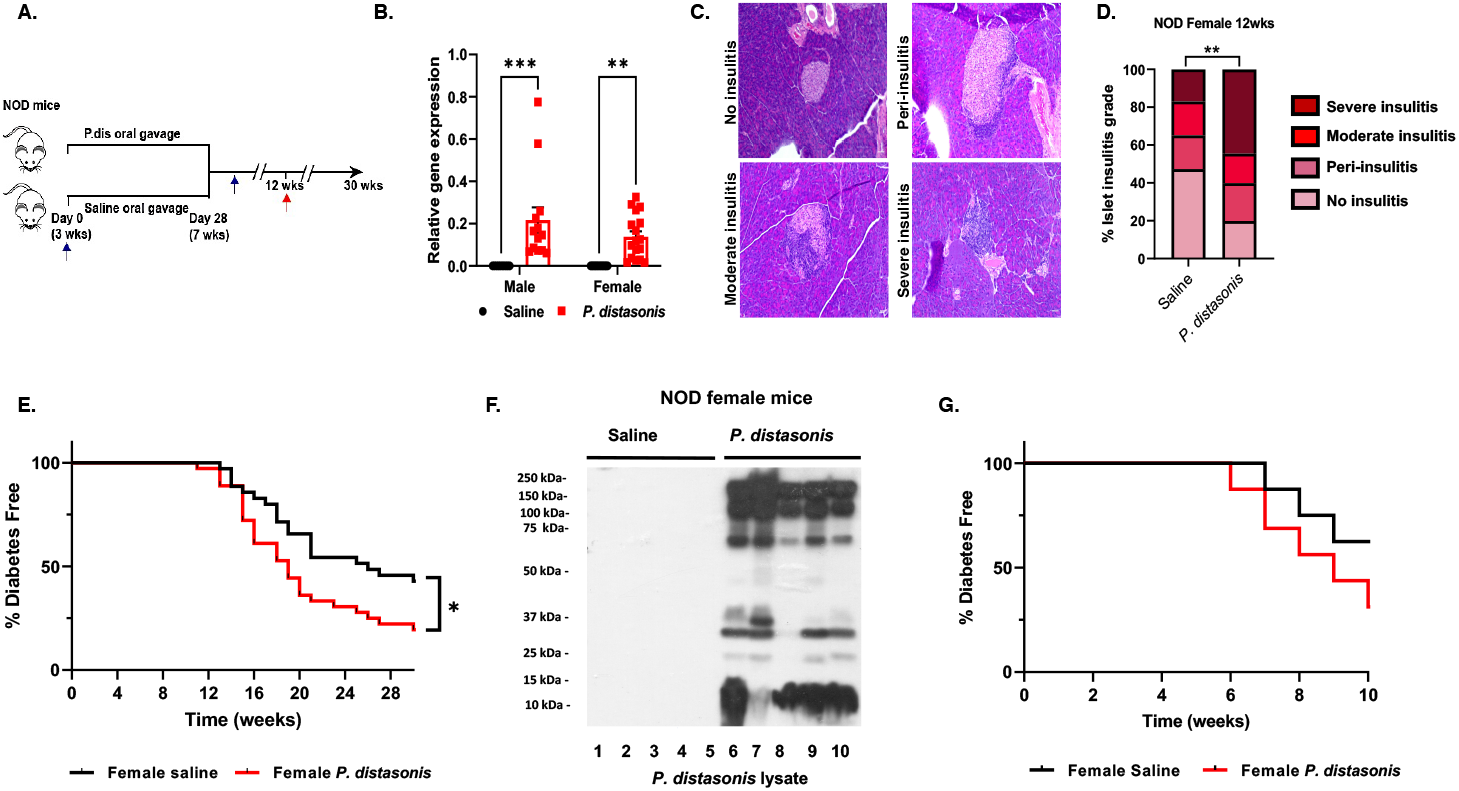
*P. distasonis* colonization enhances disease onset in female NOD mice and T1D patients have a stronger immune response to *P. distasonis* compared to healthy individuals. **(A)** Schematic overview of the *P. distasonis* oral gavage experiments (n=40/group/sex). Blue arrow shows the time-point (week 9) of fecal sample collection for qPCR experiments for *P. distasonis* colonization and red arrow shows the time-point (week 12) for pancreata collection for insulitis analysis (n = 5 mice/group/sex). **(B)** Relative abundance of *P. distasonis* in fecal samples determined by qPCR (week 12, n=10-13/male, n=12-17/female). **(C)** Representative images of islets insulitis score data where islets were scored as no insulitis, peri-insulitis, moderate insulitis or having severe infiltration as shown in images. **(D)** Quantification of insulitis scores obtained from *P. distasonis* colonized and saline gavaged female NOD mice at week 12 (n=5/group/sex). **(E)** Diabetes incidence in female NOD mice (n=35/group/sex) after daily oral gavage with either saline or *P. distasonis* for four weeks after weaning *(P<0.05)*. **(F)** Western blot analysis performed using serum samples from female NOD mice either oral gavaged with *P. distasonis* or saline (week 12, n=5/group/sex). **(G)** Diabetes incidence of the recipient NOD/SCID mice after adoptive transfer of 5×10^7^ splenocytes/mouse from individual female NOD mice to female NOD/SCID mice at 6 weeks of age (1:1 ratio, same gender, n=15). All samples in each panel are biologically independent. Log-rank (Mantel-cox) test was used for survival curves and adoptive transfer experiment. Data were expressed as mean ± SEM. *\*P<0.05*, *\*\*P<0.01*, *\*\*\*P<0.01*. Statistical analysis was performed by two-way ANOVA using Sídak multiple comparison test or two-tailed unpaired Student’s *t*-test.

To determine whether *P. distasonis* colonization stimulates an immune response against the bacterium, we performed Western blot analysis in which *P. distasonis* cellular protein lysates were resolved on SDS-PAGE, transferred to PVDF membranes, then probed using serum samples from 12-week-old *P. distasonis* colonized or control NOD mice. *P. distasonis* colonization of the NOD mice gut stimulated an antibody response to multiple bacterial proteins in both female and male NOD mice (**Figure 2F, Supplementary Figure S1C**). This response was specific; Western blots using protein lysates from *Bacteroides fragilis*, another gut microbe that has been shown to have immunomodulatory effects in NOD mice(23), produced only a weak humoral immune response in both the saline and *P. distasonis* treated groups **(Supplementary Figure S1D-E**). Importantly, serum LPS levels were not significantly different between the groups, indicating that the *P. distasonis* humoral response was not the result from of general gut barrier dysfunction stimulated by bacterial colonization **(Supplementary Figure S1F**).

To determine whether there is a similar enrichment in the humoral immune responses against *P. distasonis* occurs in humans with T1D, Western blot analysis was performed using serum samples from 12 T1D patients (median age 17.5, median disease duration 8 years, males) with age, sex and ethnicity matched controls. As shown in **Supplementary Figure S1G,** there was a very weak humoral response against *P. distasonis* in healthy subjects, strong reactivity was observed in seven out of 12 T1D patients. Thus, consistent with our findings in NOD mice, at least a subset of T1D patients appear to have a humoral immune response to *P. distasonis* proteins.

### 2.3 Adoptive transfer of splenocytes enhances T1D onset in NOD/SCID mice

To determine if the enhanced development of T1D in the NOD receiving *P. distasonis* colonized mice was T-cell mediated, we performed an adoptive transfer experiment using immunodeficient NOD/SCID mice, which lack functional B and T cells, as recipients(24). To this end, a new cohort of NOD mice was orally gavaged with *P. distasonis* or saline (n=15-20/group) as described above. At week 15, these mice were sacrificed, and 5×10^6^ splenocytes were transferred from individual NOD diabetes-free donors to 9-week-old NOD/SCID recipients (1:1 ratio, sex-matched) after which the recipients were followed for 10 weeks **(Supplementary Figure S2A**). Consistent with the effect of *P. distasonis* colonization on spontaneous T1D in NOD mice (**Figure 2G**), female NOD/SCID mice that received splenocytes from the *P. distasonis* treated group developed T1D at a higher rate than those receiving splenocytes from control NOD mice. Thus, at 10 weeks following transfer, 62.5% of recipients of receiving control splenocytes remained disease free, and this was decreased to 31% in mice receiving splenocytes from *P. distasonis* colonized NOD donors (**Figure 2G**). Hence, the immune response stimulated by *P. distasonis* in female NOD mice was sufficient to accelerate T1D in NOD/SCID mice.

### 2.5 *P. distasonis* colonization increases CD8+ T-cells and decreases Foxp3+ regulatory T cells in the splenocytes of female NOD mice

To determine the effects of *P. distasonis* treatment on NOD mice immune cell composition, a new cohort of 12-week-of age NOD mice was orally gavaged with either *P. distasonis* or saline, and the T-cell populations in splenocytes and pancreatic lymph nodes (PLNs) assessed by flow cytometry. The gating strategy for different T-cell subsets and innate immune cells are described in **Supplementary Figure S3A & B**. We found that in the spleens of *P. distasonis* colonized NOD mice there was a significant 31% increase in CD8+ T-cells, leading to a 30 % decrease in the CD4/CD8 ratio **(Figure 3A & 3B)**. PLN cells isolated from the *P. distasonis* colonized mice showed similar trends in CD4+ T-cells, CD8+ T-cells, and CD4/CD8 ratio, but these did not quite reach statistical significance **(Supplementary Figure S4A & S4B)**. Using FACS analysis, we also determined the various subsets of cells including naive cells (CD44^lo^ CD62L^hi^), effector memory cells (T_EM_, (CD44^hi^ CD62L^lo^), and, central memory cells (T_CM_, CD62L^hi^ CD44^hi^) in the TCRβ+/CD8+/CD4+ cell population in both the spleen and PLN (**Figure 3C-3F**). This revealed a significant increase in the TCRβ+/CD8+/CD4+ naive T-cell population (**Figure 3D, 3E**) and a decrease in T_EM_ CD4^+^ T-cells in splenocytes (**Figure 3C-3F**) but no alterations in TCRβ+/CD4+ naive cells in PLNs of *P. distasonis* colonized mice **(Supplementary Figure S4C-S4F**). There was also a two-fold increase in the naive cell population in both CD4+ and CD8+ T-cells in spleen (**Figure 3E & F**) and in the CD8+ naive cell population in PLNs **(Supplementary Figure S4F**). There were no differences in the central memory cells and effector memory cell populations in TCRβ+/CD8+ cells in splenocytes (**Figure 3D & 3F**) or in TCRβ+/CD8+/CD4+ cells in PLNs **(Supplementary Figure S4D-S4F**).

**Figure 3:**
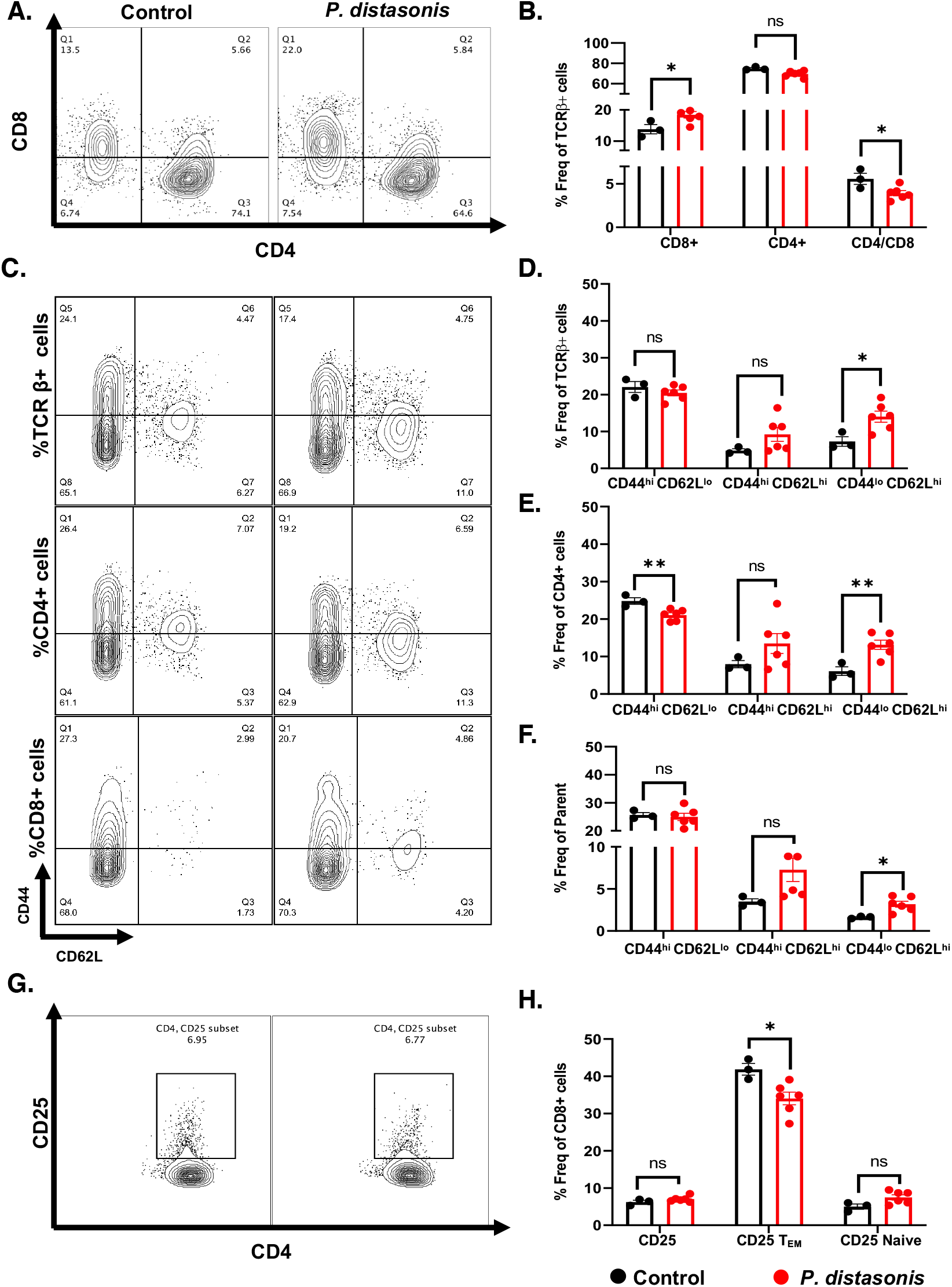
*P. distasonis* colonization increase CD8+ T-cells and Naïve cells phenotype. Mice were orally gavaged with either *P. distasonis* or saline for 4-weeks after weaning and analyzed at 12 weeks of age. **(A)** Representative image from a single experimental cohort of FACS analyses for splenic CD4+ and CD8+ T-cells from saline and *P. distasonis* gavaged mice. **(B)** CD4+ and CD8+ cells as percent of TCR-β+ immune cell subsets and ratio of splenic CD4+ to CD8+ T-cells. **(C)** CD44^lo/hi^ and CD62L^lo/hi^ T-cells in TCR-β+, CD4+ and CD8+ T-cells. **(D)** Percent of CD44^hi^ CD62L^lo^ (T_EM_), CD44^hi^ CD62L^hi^ (T_CM_), CD44^lo^ CD62L^hi^ (Naive) in TCR-β^+^ immune cell subsets, **(E)** Percent of CD44^hi^ CD62L^lo^ (T_EM_), CD44^hi^ CD62L^hi^ (T_CM_), CD44^lo^ CD62L^hi^ (Naive) in CD4+ immune T-cell subsets; **(F)** Percent of CD44^hi^ CD62L^lo^ (T_EM_), CD44^hi^ CD62L^hi^ (T_CM_), CD44^lo^ CD62L^hi^ (Naive) in CD8+ immune T-cell subsets **(G)** Representative image of FACS analyses of CD4+, CD25+ Treg population in saline and *P. distasonis* gavaged mice. **(H)** Percent of CD4+ CD25+ cells in CD4+ single cell subsets, CD44^hi^ CD62L^lo^ (T_EM_) and CD44^lo^ CD62L^hi^ (Naive) population in percent of CD4+ CD25+ single cell subsets. All samples in each panel are biologically independent. Spleens were obtained from female NOD mice oral gavaged with either *P. distasonis* (n=6) or saline (n=3). Data were expressed as means ± SEM. *, *P<0.05*, **, *P<0.01*, ***, *P<0.001*. Statistical analysis was performed by two-tailed, unpaired Student’s t-test.

T-regulatory cells (Treg) play a key role in modulating T1D autoimmunity by suppressing self-reactive T-cell proliferation(25). This is modulated by Forkhead box protein P3 (Foxp3) expression and interleukin 10 (IL-10) secretion. It has previously been shown that CD4+ CD25+ CD44+ Treg cells positively correlate with Foxp3 expression and production of IL-10(26). We found a 15 % decrease of CD4+ CD25+CD44+ Treg cells in both splenocytes (**Figure 3G & 3H**) and PLN cells **(Supplementary Figure S4G & S4H**) and a 1.6-fold increase in the total CD4+ CD25+ PLN cell population **(Supplementary Figure S4H)** of *P. distasonis* colonized mice. While there were no differences in other cell subsets **(Figure 3H).** In the CD4+ T-cell population, the percentage of Foxp3+ cells were significantly decreased in the splenocytes and PLNs of *P. distasonis* colonized NOD mice by 21% and 17.8%, respectively (**Figures 4A, 4B & S5A, representative images**). There was no difference in the pancreas and spleen weight or in the total number of the spleen and PLN cells between *P. distasonis* colonized mice and the control mice (**Supplementary Figure 5B**`). Thus, *P. distasonis* colonization decreases anti-inflammatory CD4+ CD25+ CD44+ and Foxp3+ Tregs and increase inflammatory CD8+ T-cells in splenocytes and PLNs, which could contribute to the enhancement of T1D in female NOD mice.

**Figure 4:**
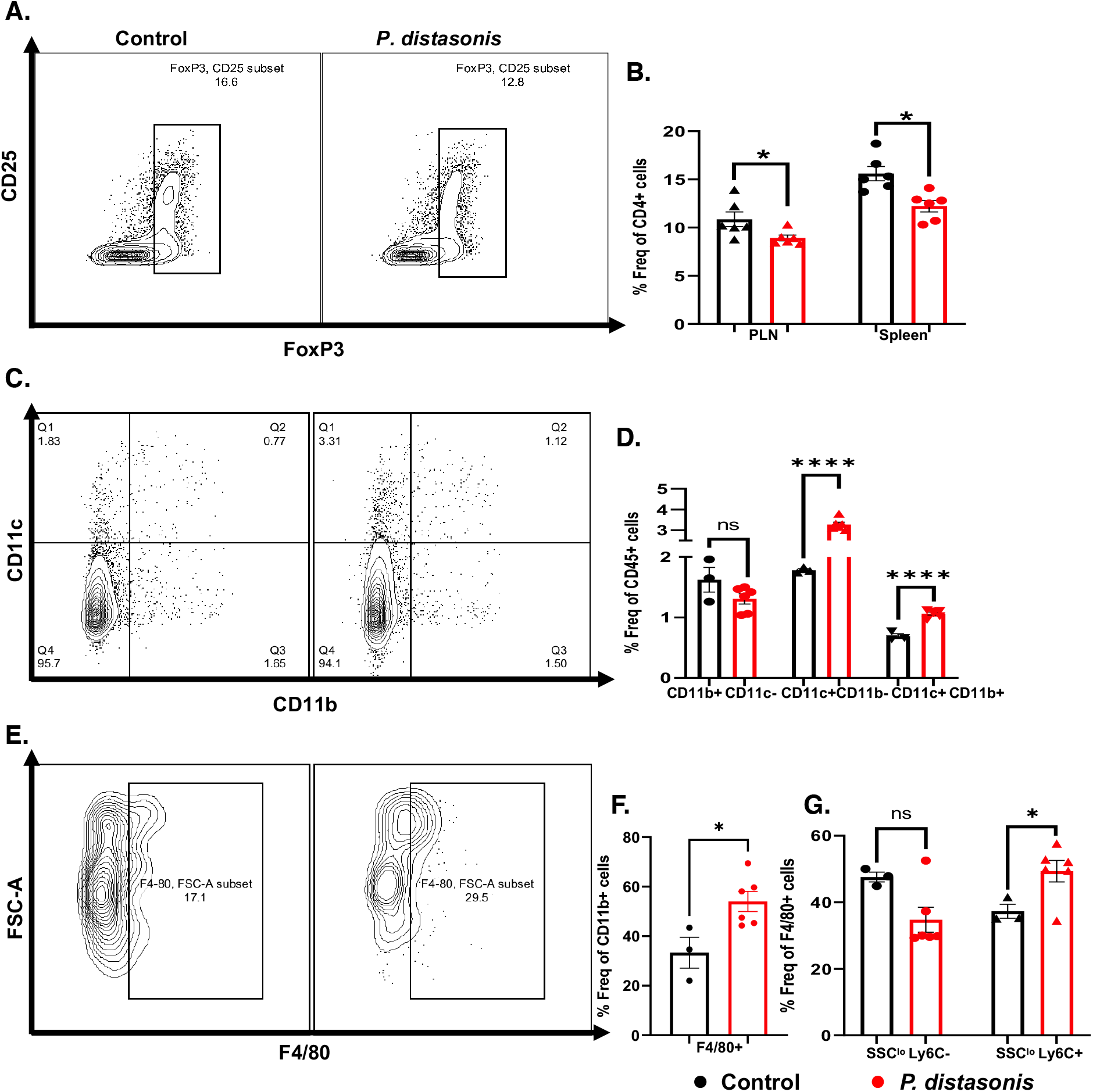
*P. distasonis* colonization decreases Foxp3+ cells and increases dendritic, and macrophage populations. **(A)** Representative image of FACS analyses of splenic Foxp3+ cells in saline and *P. distasonis* gavaged mice. **(B)** Percent of Foxp3+ cells in CD4+ T-cell subsets. **(C)** Representative image of FACS analyses of dendritic cells (CD11b-CD11c+ and CD11c+ CD11b+) in saline and *P. distasonis* gavaged mice. **(D)** Percent of CD11b+/-CD11c+/- population in splenic single cell subsets **(E)** Representative image of FACS analyses of macrophages (F4/80+cells) in spleen of saline and *P. distasonis* gavaged mice. **(F)** Percent of F4/80+ cells in CD11b+ cell subsets **(G)** Percent of F4/80+ cell subsets, where SSC^lo^Ly6C^lo^ represents residential macrophage and SSC^lo^Ly6C^hi^ represents circulatory macrophages. All samples in each panel are biologically independent. Spleens were obtained from female NOD mice oral gavaged with either *P. distasonis* (n=6) or saline (n=3). Data were expressed as means ± SEM. *, *P<0.05*, **, *P<0.01*, ***, *P<0.001*. Statistical analysis was performed by two-tailed, unpaired Student’s t-test.

### 2.6 *P. distasonis* colonization in female NOD mice increases dendritic cells and macrophages in the splenocytes

To further explore the mechanisms that lead to an increase in CD8+ T-cells in *P. distasonis* colonized NOD female mice, we assessed the number of the APCs in spleens and PLNs. CD11c+ CD11b+ dendritic cells and F4/80+ macrophages play an essential role in accelerating diabetes in NOD mice(27, 28). FACS analysis revealed a 1.5-fold increase in CD11b+ CD11c+ dendritic cells and a 1.6-fold increase in F4/80+ macrophages in the splenocytes of *P. distasonis* colonized mice (**Figure 4C-F**), with no significant differences in dendritic cells in PLNs **(Supplementary Figure S5D-S5G**). Moreover, CD11c+ CD11b-dendritic cells(29) increased 1.8-fold in the spleen of *P. distasonis* colonized mice (**Figure 4D**). There was also a 1.3-fold increase in circulatory macrophages (SSC^lo^Ly6C^hi^ cells), while there was no change in the number of residential macrophages (SSC^lo^Ly6C^lo^) in this population **(Figure 4G)**. However, there was 7-fold increase in F4/80+ macrophage population in PLN in *P. distasonis* colonized mice **(Supplementary Figure S5G**). These findings are consistent with previous studies demonstrating a role of CD11c+ CD11b+ dendritic cells and F4/80+ macrophages in accelerating diabetes in NOD mice(27, 28) and demonstrated that *P. distasonis* colonization can stimulate both innate and adaptive immune responses.

### 2.7 Critical timing of *P. distasonis hprt4-18* exposure in gut microbiome samples of children developing T1D in early life

To investigate the potential role for the *P. distasonis* insB:9-23 mimetic peptide in human T1D, we reanalyzed human gut microbiome data from the DIABIMMUNE study ^57^ using shotgun metagenomic sequencing performed on stool samples collected monthly from infants 0-3 years of age who were genetically predisposed to T1D living in Finland, Russia and Estonia and correlated this with development of islet autoantibodies, i.e., seroconversion(30). Using the metagenomic data of this study, we searched for the specific DNA sequences encoding the *P. distasonis hprt4-18* peptide. We found that in the children who remained negative for autoantibody markers for T1D, sequences encoding *P. distasonis hprt4-18* were detectable in 20-40% patients with no significant change between ages 0 to 3. By comparison, in children who later became autoantibody positive, *hprt4-18* encoding sequences were absent in the first year of life, began to appear by age 2 and then were found in 60% of the children by age 3 (**Figure 5A**).

**Figure 5.**
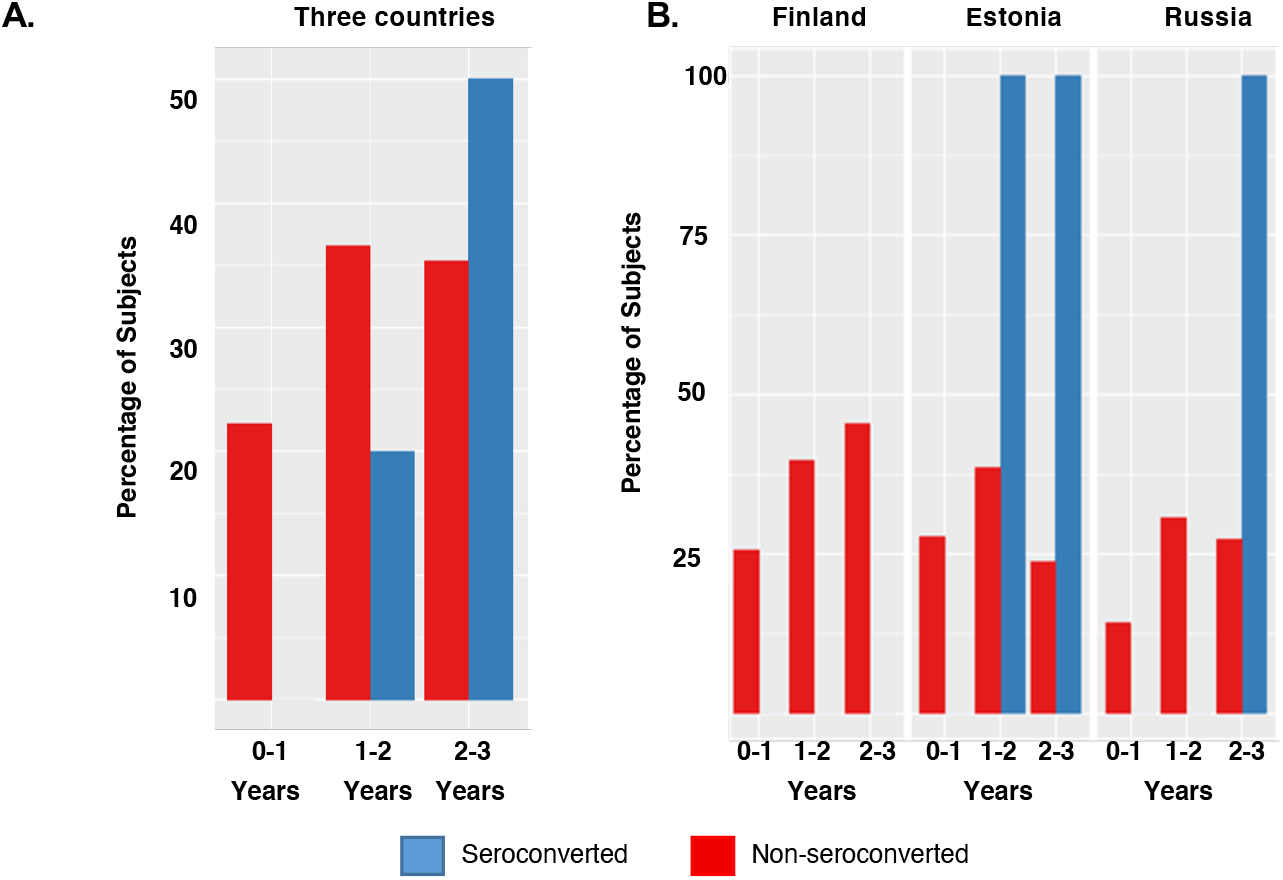
Reanalysis of DIABIMMUNE gut microbiome data revealed the presence of sequences encoding *P. distasonis hprt4-18* peptide in all children developing autoantibodies in Estonia after age 1 and Russia after age 2. **(A-B)** Reanalysis of DIABIMMUNE gut microbiome data for the presence of *hprt4-18* peptide **(A)** The percentage of subjects having sequence encoding *hprt4-18* in the gut in all three countries (Finland, Estonia and Russia) and **(B)** in individual countries at different phases of gut microbiota development (age 0 to 3). Blue represents subjects that developed autoantibodies (seroconverted) and red represents subjects with no autoantibodies (non-seroconverted). 16 of 74 subjects in Finland, 14 of 74 of subjects in Estonia, 4 of 74 subjects in Russia developed autoantibodies during the study.

The exact frequency and timing of appearance of the *hprt4-18* sequence in the microbiome varied by country. Thus, in subjects from Estonia (14 of 74 infants) and Russia (4 of 74 infants), for those who became seropositive, we did not find the *hprt4-18* sequence in their gut microbiota during the first year of life, but was found in 100% of subjects who developed autoantibodies by the age of 2 or 3 (**Figure 5B**). In contrast, this sequence was found in 15-40% of the seronegative infants throughout this time period. Interestingly, while children in Finland who were seronegative has similar prevalence of the microbial sequence as those in Estonia and Russia, infants who became seropositive in Finland (16 of 74 infants) had no detectable *hprt4-18* sequences during the first three years of life (**Figure 5B**). These results suggest that a delayed exposure to the *P. distasonis hprt4-18* peptide, i.e., after the first year of life, may be able to trigger or potentiate an autoimmune response that leads to development of T1D in genetically susceptible individuals. This is also consistent with our findings in NOD mice where colonization of the female NOD mice with *P. distasonis* after weaning accelerated diabetes development.

## 3. DISCUSSION

Despite major increases in our understanding of the role of autoimmunity in pathogenesis of T1D, the triggering events which lead to disease development remain poorly understood. It is well known that an important component of the early autoimmune response in individuals who ultimately develop T1D is the development of autoantibodies and T-cell reactivity to islet proteins, especially insulin. While there are multiple potential epitopes, the dominant sequence within the insulin molecule to which reactivity occurs is a sequence in the B-chain involving amino acids 9-23(14). This B:9-23 peptide (SHLVEALYLVCGERG) can bind to DQ8 molecules utilizing three different registers, with the amino acid V at position 12, E at position 13 or A at position 14(31). It has been shown that a RàE substitution at position 22 of the B:9-23 peptide creates an even more potent agonist for activating B:9-23 T cell clones, implicating the A at position 14 as the p1 anchor(18).

In this study, we have investigated the potential role of molecular mimicry as a link between microbial flora and this component of T1D development. Molecular mimicry mechanisms are based on the degeneracy of T-cell recognition(32, 33) and can be either pathogenic(34) or protective(35). While molecular mimicry has long been postulated as a potential factor in autoimmune diseases, including T1D (36, 37), progress in this area has been limited due to a lack of identification of microbial sequences which might trigger this response(38). In the current study, taking advantage of the growing genome databases for microbes in the environment, we have identified 47 microbial peptides with high sequence homology to insB:9-23. We demonstrate that of these, a peptide with the sequence of *hprt4-18* from *P. distasonis,* can be recognized by and activate both murine and human T-cells known to respond to insB:9-23 and can cross-stimulate an immune response to the insB:9-23 peptide in mice. Moreover, exposure of NOD mice to *P. distasonis* in the gut microbiome during early life results in increased insulitis and accelerates diabetes onset. This pathogenic effect is mediated by CD8+ T splenocytes, as evidenced by the ability to adoptively transfer this increase in diabetes risk to female NOD/SCID mice.

Microbiota and gut microbiota in particular have been shown to play a role in modulating T1D onset in NOD mice(7, 39–41), but previous reports have identified primarily protective effects, and most through somewhat non-specific mechanisms. For example, NOD mice reared in germ-free environments have higher rates of development of diabetes than those raised in conventional facilities(40). In terms of protective effects of some specific bacterial species, the presence of segmented filamentous bacteria in the gut correlates with diabetes protection in NOD female mice, which normally have a high incidence of disease(42). Likewise, female NOD mice colonized at 3-10 weeks of age with *Akkermensia muciniphila* show delayed diabetes development(43), and oral administration of heat-killed *B. fragilis* has been shown to suppress autoimmunity in NOD mice when administered under conditions which induce increased gut permeability(23). Since literally hundreds of treatments have been shown to decrease development of diabetes in NOD mice (44, 45), however, the specificity of these effects has been questioned(46). By contrast, in the present study we show that exposure to *P. distasonis* can accelerate disease development in NOD mice and that a specific peptide encoded in their genome, *hprt4-18*, can serve as a mimic of the major insulin auto-epitope at position B9-23. This peptide and the insulin peptide can also stimulate a bidirectional cross-reactive immune response, proving its nature as a molecular mimic.

In addition to the immunologic cross-reactivity, our study also provides some insight into the mechanisms of immune cell regulation by this bacterium. Thus, colonization of NOD mice with *P. distasonis* increases the CD8+ T-cell population and decreases the CD4/CD8 T-cell ratio. A reduction in this ratio has been previously shown to be associated with increased T1D in humans(47). G9C8 CD8+ T-cell clones originally isolated from the islets of young NOD mice cells are activated by an insB:15-23 peptide, i.e., the C-terminal fragment of the insB:9-23 peptide, and this accelerates diabetes when adoptively transferred in NOD/SCID mice, even in the absence of CD4+ T-cells(48, 49). Wong *et al.* showed that cross presentation of insulin by dendritic cells can stimulate G9C8 T-cell clones, and both insulin and the insB:15-23 peptide can stimulate proliferation of the naive cell phenotype in NOD mice (48). The *P. distasonis* sequence is identical to this peptide in 6 of its 9 residues. We also observed that *P. distasonis* colonization in NOD mice decreased Foxp3+ Treg cells in spleen and PLNs. This effect on Foxp3+ Treg cells is particularly interesting because Foxp3+ Treg cells are dysregulated in newly diagnosed T1D patients(50) and increasing Foxp3+ Treg cells in NOD mice can delay the onset of diabetes in NOD mice(25, 51).

A plethora of human gut microbiome studies have demonstrated that the composition of gut microbiota in patients with autoimmune diseases, including multiple sclerosis(52), systemic lupus erythematosus, anti-phospholipid syndrome(53), Crohn’s disease(54–56), ulcerative colitis(57), inflammatory bowel diseases(58, 59), and celiac disease(60) are significantly different from those in healthy controls. Although the methodologies and conclusions differ in studies of microbiota from subjects with or at-risk for T1D(8, 30, 61), most show that the diversity of gut microbiota is decreased in T1D patients with increased prevalence of *Bacteroidetes* species. This is also associated with an altered serum metabolomic profile compared to healthy controls(7). While most of these studies have not been able to define any mechanism by which this might relate to disease development, data in the DIABIMMUNE study^57^ has suggested that the higher T1D rates in Finnish Karelia and Estonia compared to Russian Karelia may in part be related to an action of different lipopolysaccharides (LPS) present in the gut microbiota on the immune response.

The Environmental Determinants of Diabetes in the Young (TEDDY) study, which has collected over 12,000 fecal samples from 903 children at risk for T1D in four countries, has identified *Parabacteroides* as the genus most significantly associated with T1D(62). This is consistent with our findings that a *P. distasonis* peptide can induce T-cells and antibodies that cross-react with the major epitope in insulin involved in the autoimmune response. Reanalysis of the DIABIMMUNE metagenomic data show that 100% of the children becoming seropositive for autoantibodies related to T1D in Russia and Estonia have the DNA encoding the *P. distasonis* hprt4-18 peptide in their gut microbiota in first 2-3 years of life compared to less than 40% of children remaining seronegative. Interestingly, most of these seroconverters were negative for this microbial sequence in the first year of life, suggesting the timing of exposure to this microbial antigen may be important in disease development. The TEDDY study showed that human gut microbiota development can be divided into three phases: a developmental phase (months 3-14), a transitional phase (months 15-30), and a stable phase (months 31-46)(62). Our data suggest that exposure to *P. distasonis* in the transitional phase may be critical in T1D pathogenesis. Further studies will be needed to directly investigate the link between human T1D and *P. distasonis*, but the cross-reactivity of the *P. distasonis hprt*4-18 peptide with insB:9-23 activated human T cell clones represents a clear potential mechanism.

While we have focused only on insulin and the B:9-23 peptide as an antigenic determinant of disease, there are clearly other epitopes recognized by both circulating and islet-infiltrating T-cells in the insulin B-chain(63), A-chain(64, 65), or C-peptides(16, 66). The recent discovery that hybrid insulin peptides (HIPs) may be novel epitopes formed by post-translational modifications formed in the beta cell(67, 68) also opens the possibility of microbial sequences that may more closely mimic these hybrid peptides. Defective ribosomal products (Drips) of insulin have also been shown to lead to aberrant insulin polypeptides rendering beta cells immunogenic(69). We must also keep in mind that insulin is not the only antigen to which antibodies or reactive T-cell lines are formed in T1D. Other islet antigens include GAD, IA-2, and HIPs(70). Given the enormous number of microbial peptides produced by the gut microbiota, we expect that other microbial peptides may exist with the potential to mimic these epitopes and/or trigger related autoantigen reactive T-cells. Indeed, Tai *et al.* identified a microbial peptide mimic produced by *Fusobacteria* that can stimulate islet-specific glucose-6-phosphatase (IGRP) specific mouse T-cells and promote diabetes development in a new Toll-like receptor (TLR) deficient (TLR^−/−^) and MyD88^−/−^ NY8.3 NOD mouse model(71).

In summary, our data define a novel molecular mimicry mechanism in which a specific sequence in a normal commensal gut microbe can mimic a sequence in insulin B-chain and trigger or modify the immune response involved in development of T1D. This finding may provide a new target for treat and a window of opportunity to prevent or delay T1D development. These data also have implications for other diseases with an autoimmune component. Today, we have databases with enormous amounts of microbial and microbiome sequence data, which can be leveraged to address the role of molecular mimicry in the autoimmunity, not only of T1D, but also lupus erythematosis(72), inflammatory cardiomyopathy(73), and multiple sclerosis(74). Our findings demonstrate a new link to gut antigens and autoimmune diseases with the potential to ultimately provide new tools, including vaccines, antibiotics, or probiotics for the prevention and treatment of autoimmune diseases.

## 4. MATERIAL AND METHODS

### 4.1 Bioinformatics

A bioinformatics search was performed using NCBI BLASTp for the presence of the microbial peptide sequences with homology to human insB:9-23 sequence as query against viral (NCBI taxonomic ID: 10239), bacterial (NCBI taxonomic ID: 2), and fungal (NCBI taxonomic ID: 10239) proteomes. The whole peptide sequence of each significant hit was compared with insB:9-23 using a multiple sequence alignment program (Clustal Omega) to determine the number of identical and conserved residues. This yielded the data in **Supplementary Table S1**. An additional bioinformatics search was performed to determine whether *P. distasonis* peptide is unique. *P. distasonis* peptide (15aa) or *P. distasonis* hypoxanthine phosphoribosyltransferase *(hprt4-18)* protein sequences were used as queries and searched against BLASTp using non-redundant protein sequences.

### 4.2 Bacterial culture

*Parabacteroides distasonis* D13 strain was purchased from Dr. Emma Allen-Vercoe’s laboratory at the University of Guelph. Dr. Allen-Vercoe’s group isolated this bacterium from the colon of an ulcerative colitis patient. *P. distasonis* D13 strain and *Bacteroides fragilis* (ATCC^®^ 25285) were cultured in anaerobic culture broth (Tryptic Soy Broth supplemented with 5 µg/ml Hemin (BD Biosciences) and 1µg/ml Vitamin K1 (VWR) at 37°C in an anaerobic chamber (Coy Laboratory Product).

### 4.3 Animals

NOD/ShiLtJ (NOD) and NOD. Cg-*Pcrkdc scid*/J (NOD/SCID) mice were purchased from Jackson Laboratory. Mice were maintained and bred in the Boston College Animal Care Facility.

The mice were maintained under specific pathogen-free conditions in a 12-hrs dark/light cycle with autoclaved food, water, and bedding. All experiments complied with regulations and ethics guidelines of the National Institute of Health and were approved by the IACUC of Boston College (Protocol No.#B2019-003 and 2019-004). For colonization, *P. distasonis* bacteria were re-suspended in saline at a concentration of 10^9^ CFU/mL and oral gavaged using 22ga plastic feeding tube (Instech Laboratories) (100μl/mouse, 10^8^ CFU/mouse). 3-week-old NOD mice were orally gavaged with *P. distasonis* D13 daily for four weeks right after weaning for 4 weeks. Control groups were gavaged with sterile saline.

To determine colonization, the DNA was extracted from 100 mg mouse fecal samples (9-week-old, 2 weeks after final oral gavage treatment) using QIAamp PowerFecal Pro DNA kit (Qiagen) following the manufacturer’s instruction. qPCR was conducted using QuantStudio 3 Real-Time PCR System (Applied Biosystems) with Power SYBR Green Master Mix (Applied Biosystems) following manufacturer’s instruction. The primers were described previously and listed in Supplementary **Table S2**(75, 76). The relevant abundance of *P. distasonis* was determined through being normalized with *Eubacteria*.

### 4.4 Human T-Cell Clone Stimulation

Twenty 15-mer peptides **(Figure 1A**) used in human and mouse T-cell stimulation experiments were chemically synthesized by Genscript (TFA removal, >85% purity). The peptides were dissolved in DMSO at 20 mg/mL (∼12 mM). Human T-cell clone stimulation assays were performed as previously described(77). Briefly, insB:11-23-specific T-cell clones were stimulated in 96-well round-bottom plates with irrelevant, specific, or microbial mimotope peptides in the presence of irradiated DQ8cis- or DQ8trans-expressing HEK293 cells as APCs. 50 μl of supernatants from cultures of T cell clones were collected after 48 hrs of stimulation and added to each well of 96-well round-bottom plates precoated with IFN-γ (clone MD-1) capturing antibodies (BioLegend). After overnight incubation, bound cytokines were detected by biotinylated anti-IFN-γ (clone 4s.B3) and quantified using a Victor2 D time-resolved fluorometer (Perkin Elmer). The % activity was calculated by dividing the SI of mutated peptides by the SI of the wild-type peptide. Experiments were performed in the presence of 1 μg/ml of anti-CD28 antibodies. Unless otherwise stated, peptide concentrations were 2.5 μM.

### 4.5 NOD mice T-cell stimulation and antigen presentation assay

This experiment was performed as described previously(22). Briefly, the peptides were ordered from Genscript (TFA removal, >85% purity) and dissolved in a base buffer consisting of 50 mM NaCl, 10 mM HEPES, pH 6.8 with 200 µM TCEP. *P. distasonis hprt4-18* peptide - (**RILVELLYLVCSEYL)** was dissolved in 5% DMSO (the final DMSO concentration is less than 1%). The C3g7 cell line, which expresses an abundance of MHC-II I-A^g7^, was used as APCs and was treated with 1/2log dilutions of peptide (starting from 10 µM). These cells were then cultured with the IIT-3 T-cell hybridomas for 18 hrs. Culture supernatants were assayed for IL-2 by incubation with the CTLL-2 cell line. CTLL-2 cells are responsive to IL-2 and only actively divide in the presence of IL-2. Next, CTLL cell proliferation was determined by 3H uptake (CPM) using a scintillation counter.

### 4.6 Immunization of the NOD mice

The experiments were performed as described previously(78). Briefly, 13 weeks old male NOD mice (n=3 mice per group) were immunized in the footpad with the either *hprt4-18* peptide or insB:9-23 peptide (10 nmol/mouse). After 7 days, the draining (popliteal) lymph nodes were removed and pooled for examination by ELISpot. In the ELISpot assay, node cells were recalled with the various peptides to elicit either an IL-2 or IFN-γ response. Spots were analyzed by Immunospot 5.0 (C.T.L.)

### 4.7 Reanalysis of the published metagenomics data

DIABIMMUNE data was downloaded with the help of Dr. Alex Kostic. The reads were assembled using SPAdes as the reads were not ideal for peptide search(30) and are now available at https://github.com/ablab/spades. These assembled contigs were used for a tBLASTn search to identify “RILVELLYLVCSEYL” encoding sequences in these samples. The data was classified based on the country and collection time of the stool samples, e.g., 0-1, 1-2, and 2-3 years.

### 4.8 Splenocyte transplantation

Splenocytes were isolated from 15 weeks donor NOD mice by filtering through a sterile 70 µm nylon mesh. The isolated splenocytes were incubated for 3 mins in red blood cell lysis with ACK (Lonza) followed by washing with media. The isolated splenocytes were intravenously injected (5×10^7^ cells/mouse) through lateral tail vein into recipient NOD/SCID mice (9 weeks old). Donor NOD mice and recipient NOD/SCID mice were sex-matched. Recipient mice were monitored for diabetes by checking tail vein blood glucose levels (>250 mg/dl) twice per week. The experiment terminated 10 weeks after the transplantation or unless the mice were diagnosed with diabetes. Donor mice were 15-week male or female NOD mice colonized with either *P. distasonis* or their sex-matched saline control (n=15-19).

### 4.9 FACS and Flow Cytometry

We collected spleen and Pancreatic Lymph Nodes (PLNs) cells from saline-treated and *P. distasonis* colonized female NOD mice at 12 weeks age. Single-cell suspension was obtained by mechanical disruption of spleen and PLNs using an Ammonium-Chloride-Potassium (ACK) lysis buffer for 5 minutes followed by filtration with a 70-µm filter. For staining of surface markers, cells were incubated in fluorescently labeled antibodies **(Supplementary Table S4**) for 15 minutes in PBS containing 2% FBS at room temperature. Cells were then washed with 2 mL of staining buffer and fixed with 1% Paraformaldehyde (PFA). For Foxp3 staining, fix and perm permeabilization kit (Thermo fisher) was used as per manufacturer instruction. All the flow cytometry antibodies were obtained from BioLegend, and Flow cytometry was performed using BD FACSAria III sorter (BD biosciences) at the Boston College Core facility. The data was analyzed using FlowJo 10.0 software. Gating strategy for the determination of different cells subsets are described in supplementary **Figure S2.**

### 4.10 Insulitis score

Pancreata were isolated from *P. distasonis* colonized 12 weeks NOD mice and control mice (n=5) followed by fixation with 4% paraformaldehyde overnight at 4°C. Pancreata were embedded in paraffin blocks and were sectioned into 5µm thickness. To determine insulitis scores, pancreatic sections were stained with hematoxylin-eosin at Harvard University BIDMC Histology Core. Slides were analyzed using EVOS XL Core microscope with an LPlan PH2 10 × /0.25 and an LPlan PH2 20×/0.40 objective. The pancreatic insulitis was screened and evaluated as described previously(79).

### 4.11 Western Blot analysis

Protein lysate was extracted using CelLytic B Cell Lysis Reagent (Sigma) following the manufacturer’s instruction. Protein concentrations were determined using BCA assay (ThermoFisher Scientific). 20 µg of total denatured bacterial protein was loaded into each well in SDS gel. Wet transfer was performed by using PVDF membrane for 1 h at 4°C. The membrane was then blocked with 5% non-fat milk in PBST. After washing, the membrane was incubated with mouse serum (1: 2,500) or human serum (1: 10,000) overnight at 4°C followed by washing three-time wash with PBST and incubation with secondary antibodies, sheep anti-mouse IgG antibody (1:3000, Millipore) or goat anti-human IgG antibody (1:3000, SantaCruz) for 1 h at room temperature. The membrane was washed and developed with chemiluminescent substrate ECL (ThermoFisher Scientific).

### 4.12 Endotoxin ELISA

12-weeks old NOD mice serum was collected and stored at −80 °C until use. Serum endotoxin levels were measured using a Pierce Chromogenic Endotoxin Quant Kit (ThermoFisher Scientific) following the manufacturers’ instructions.

### 4.13. Human Plasma Samples

Peripheral blood was collected from living subjects following the provision of written informed consent (and assent in the case of minors) in accordance with IRB approved protocols (University of Florida IRB# 201400703) and the Declaration of Helsinki. The patient demographics outlined in **Supplementary Table 3**.

### 4.14 Statistics

Data are presented as the mean ± SEM. Survival curves were analyzed by a log-rank (Mantel-cox) test. Statistical significance was evaluated using unpaired two-tailed Student’s *t*-test for two-group comparison or ANOVA, followed by Dunnet’s or Šidák’s multiple comparisons tests whichever was recommended. A *p-value* of less than *0.05* was considered significant (*) at the confidence interval of 95%.

## Acknowledgements

We first want to thank Prof. Emil Unanue and Dr. Anthony Vomund (University of Washington, St. Louis) for testing microbial insulins on T-cell hybridomas, immunization experiments, and their help to interpret the data. Thanks to Jonathan Dreyfuss and Hui Pan (Joslin Bioinformatics Core) for bioinformatics analysis. Thanks to Babak Momeni (Boston College) for sharing his laboratory’s anaerobic chamber. We thank Bret Judson and the Boston College Imaging Core for infrastructure and support. We acknowledge our undergraduate students Tu Tran, Ruixu Huang, David Kim, Scott Hsu, Kaan Sevgi and Maximilian Figura and Typhania Zanou for their help with the microscopy work and the animal experiments. Thanks to Nancy McGilloway and Todd Gaines for their support in the Boston College Animal Care Facility. We thank Alex Kostic, Tao Xu, Thomas Serwold, and Shio Kobayashi (Joslin Diabetes Center) for valuable Discussion.

## Author contributions

K.G assisted with NOD mice experiments, NOD mice colonization followed by FACS staining, data analysis and writing the manuscript. Q.H assisted with NOD mice experiments; NOD mice oral gavage, splenocytes transfer, qPCR, and Western blot analysis using mice serum. I.C and W.K assisted with the experiments using NOD mice T-cell clones. C.B and A.R assisted with oral gavage experiments and following the diabetes onset in NOD mice. P.A assisted with FACS facilities and instrument handling. CRK assisted with research design and data analysis. All authors assisted with the analysis of the data. E.A completed bioinformatics analysis for the mimic identification, supervised the project, analyzed the data and wrote the paper.

## Data and materials availability

Because the DIABIMMUE study metagenomics sequencing reads were not ideal for a peptide search, we first assembled the reads using SPAdes (11), and they are available at https://github.com/ablab/spades.

