## Supplemental document for "A Gut Microbial Peptide and Molecular Mimicry in the Pathogenesis of Type 1 Diabetes"

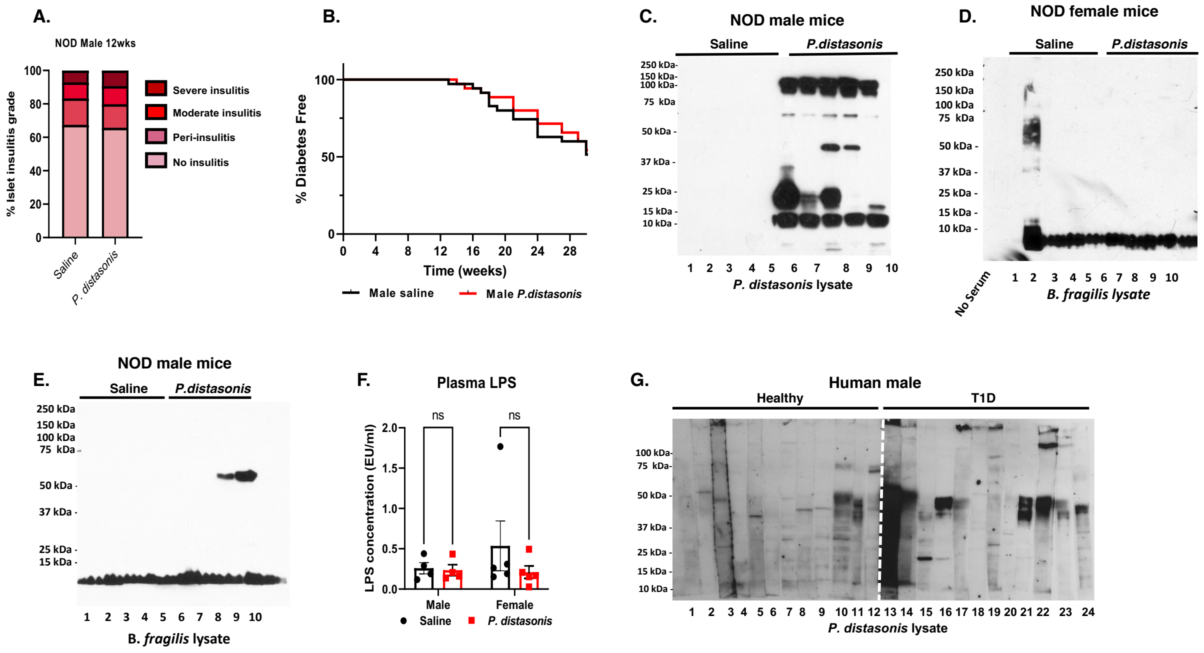


**Figure S1. Effects of *P. distasonis* colonization in mice and humans. (A)** Quantification of insulitis scores obtained from *P. distasonis* colonized and saline gavaged male NOD mice at week 12 (n=5/group/sex). **(B)** Diabetes incidence in NOD male mice (n=35/group/sex) after daily oral gavage of with either saline or *P. distasonis* for four weeks after weaning. **(C)** Western blot analysis against *P. distasonis* lysate using serum samples from male NOD mice orally gavaged with either *P. distasonis* or saline (week 12, n=5/group/sex). **(D-E)** Western blot analysis against *B. fragilis* lysate using plasma samples from (**D**) female and (**E**) male mice, orally gavaged with either *P. distasonis* or saline (week 12, n=5/group/sex). **(F)** Endotoxin levels in NOD mice serum (week 12, n=5/group/sex). **(G)** Western blot analysis against *P. distasonis* lysate using plasma samples obtained from T1D patients or healthy subjects (n=12/group). All samples in each panel are biologically independent.

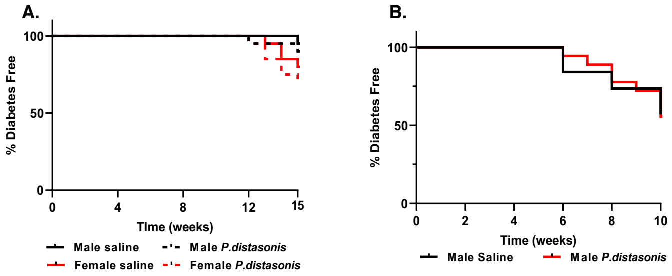


**Figure S2. Adoptive transfer of splenocytes to male NOD/SCID mice. (A)** Incidence of diabetes in donor NOD mice until adoptive transfer of splenocyte (n=16-20/group/sex). **(B)** Diabetes incidence of the recipient NOD/SCID mice after adoptive transfer of $5x10^7$ splenocytes/mouse from individual NOD male mice to NOD/SCID male mice at 6 weeks of age (1:1 ratio, same gender, n=17-19).

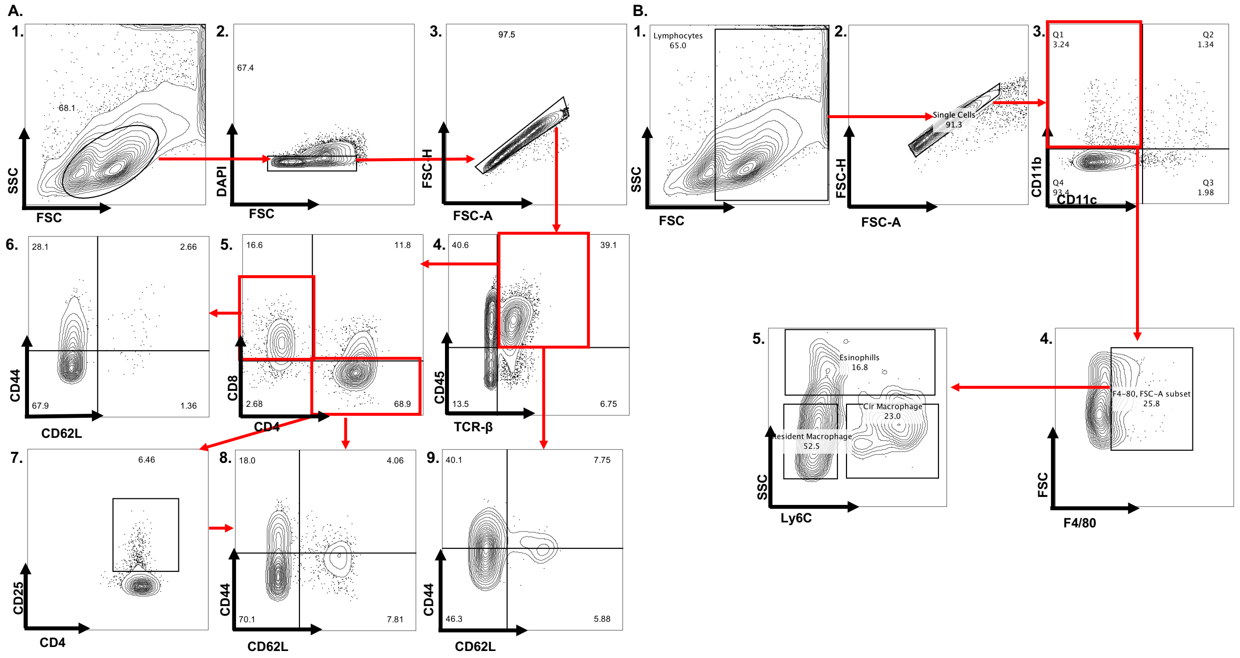


**Figure S3: The gating strategy for flow cytometry assay. (A)** Gating strategy for T-cell subsets. **(1)** FSC vs SSC gating to discriminate mononuclear cells and polymorphonuclear cells. **(2)** DAPI gating for live cells. **(3)** FSC-H vs FSC-A for singlet cell selection. **(4)**

CD45+TCRβ+was used for the selection of T cells. **(5)** CD4 vs CD8 for helper and cytotoxic T-cells, (**7**) CD4 vs CD25 for CD4+CD25+ cells (**6, 8, and 9**) CD44 vs CD62L to determine naïve, effector memory and central memory cells in CD8+, CD4+, TCRβ+ and CD4+CD25+ cell subsets. **(B)** Gating strategy for innate immune cells. **(1)** FSC vs SSC gating to discriminate mononuclear cells and polymorphonuclear cells. **(2)** FSC-H vs FSC-A for singlet cell selection. **(3)** CD11b vs CD11c cells to determine CD11b+, CD11c+ and Cd11b+Cd11c+ cell subsets. **(4)** Cd11b+ cells were selected for F4/80+ macrophages. **(5)** SSC vs Ly6C to determine circulatory and residential macrophages.

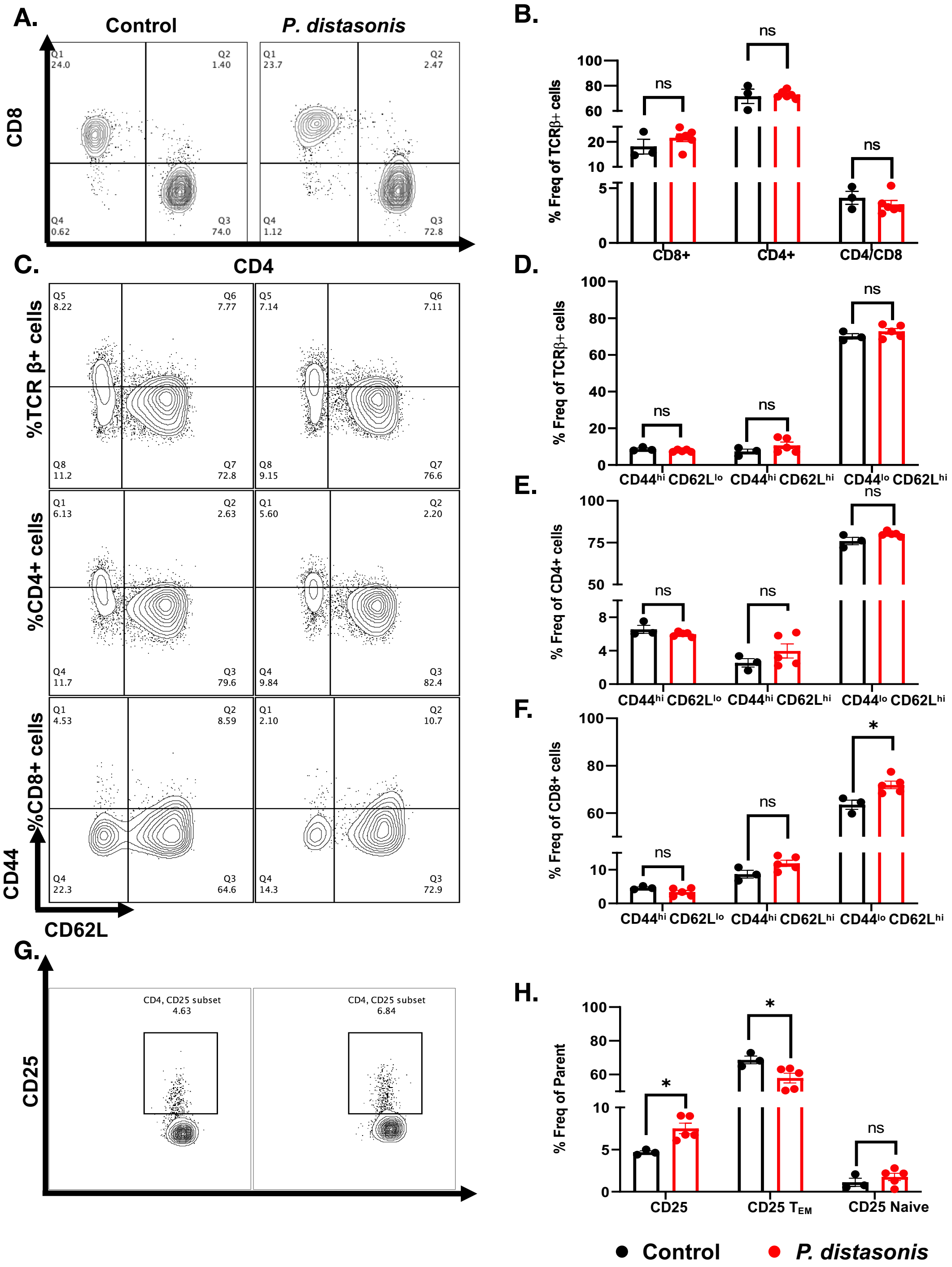

A.
Control
P. distasonis
CD8
CD4
B.
% Freq of TCRβ+ cells
ns
CD8+
CD4+
CD4/CD8
C.
%TCR β+ cells
%CD4+ cells
%CD8+ cells
CD44
CD62L
D.
CD44hi CD62Llo
CD44hi CD62Lhi
CD44lo CD62Lhi
E.
% Freq of CD4+ cells
F.
% Freq of CD8+ cells
*
G.
CD25
CD4, CD25 subset
4.63
6.84
H.
% Freq of Parent
CD25
CD25 TEM
CD25 Naive
Control
P. distasonis

**Figure S4: The effects of *P. distasonis* colonization on pancreatic lymph nodes (PLNs) immune cell composition**. Mice were orally gavaged with either *P. distasonis* or saline for four weeks after weaning and analyzed at 12 weeks of age. **(A)** Representative image from a single experimental cohort of FACS analyses for PLNs CD4+ and CD8+ T-cells from saline and *P. distasonis* gavaged mice. **(B)** CD4+ and CD8+ cells as percent of TCR-β+ immune cell subsets and ratio of PLNs CD4+ to CD8+ T-cells. **(C)** $CD44^{lo/hi}$ and $CD62L^{lo/hi}$ T-cells in TCR-β+, CD4+ and CD8+ T-cells. **(D)** Percent of $CD44^{hi}$ $CD62L^{lo}$ ($T_{EM}$), $CD44^{hi}$ $CD62L^{hi}$ ($T_{CM}$), $CD44^{lo}$ $CD62L^{hi}$ (Naive) in TCR-β+ immune cell subsets, **(E)** Percent of $CD44^{hi}$ $CD62L^{lo}$ ($T_{EM}$), $CD44^{hi}$ $CD62L^{hi}$ ($T_{CM}$), $CD44^{lo}$ $CD62L^{hi}$ (Naive) in CD4+ immune T-cell subsets; **(F)** Percent of $CD44^{hi}$ $CD62L^{lo}$ ($T_{EM}$), $CD44^{hi}$ $CD62L^{hi}$ ($T_{CM}$), $CD44^{lo}$ $CD62L^{hi}$ (Naive) in CD8+ immune T-cell subsets **(G)** Representative image of FACS analyses of CD4+, CD25+ Treg population in saline and *P. distasonis* gavaged mice. **(H)** Percent of CD4+ CD25+ cells in CD4+ single cell subsets, $CD44^{hi}$ $CD62L^{lo}$ ($T_{EM}$) and $CD44^{lo}$ $CD62L^{hi}$ (Naive) population in percent of CD4+ CD25+ single cell subsets. All samples in each panel are biologically independent. PLNs were obtained from female NOD mice orally gavaged with either *P. distasonis* (n=6) or saline (n=3). Data were expressed as means ± SEM. *, $P<0.05$, **, $P<0.01$, ***, $P<0.001$. Statistical analysis was performed by two-tailed, unpaired Student's t-test.

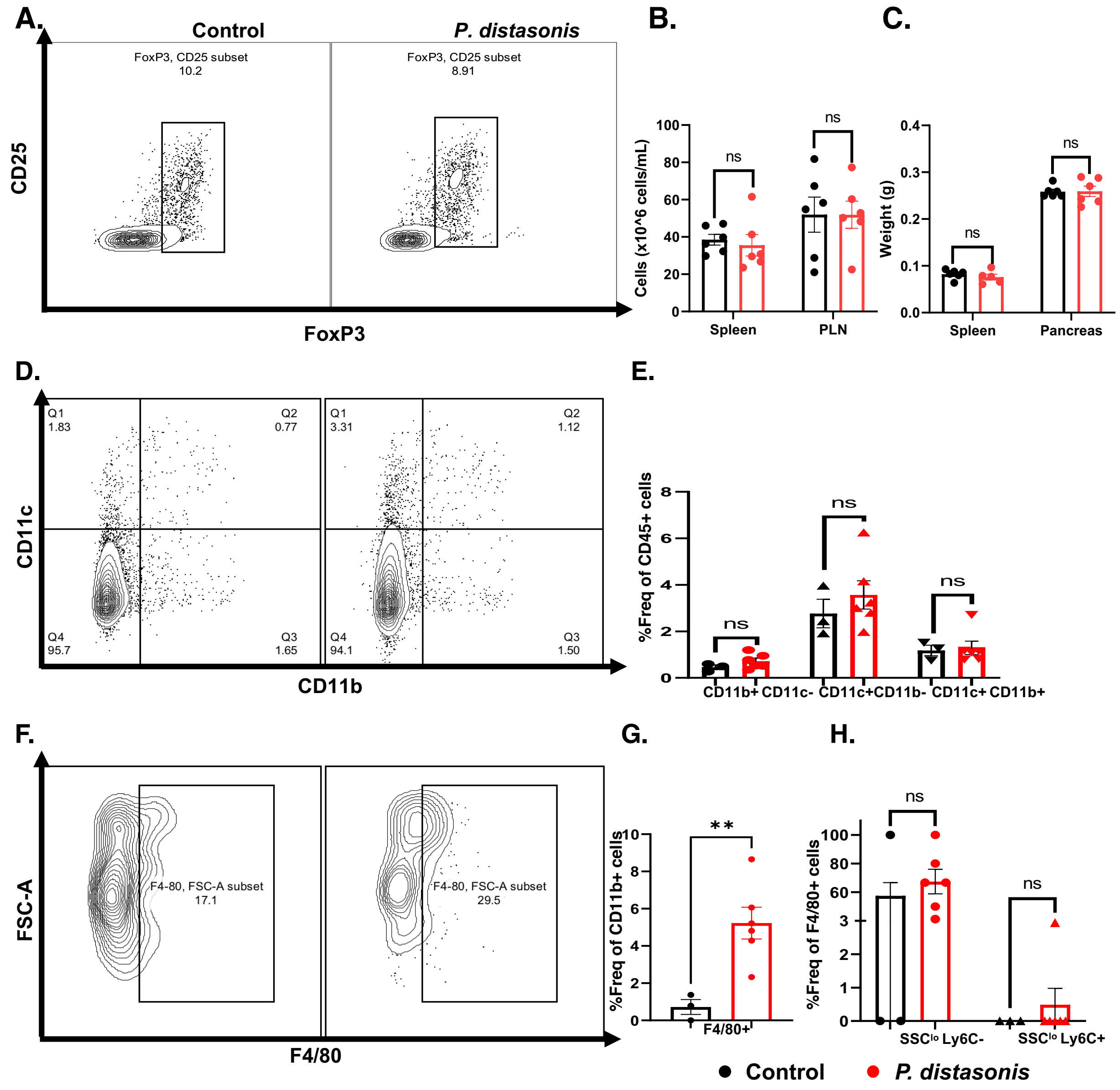


**Figure S5:** ***P. distasonis*** **colonization increase macrophage population in pancreatic lymph nodes (PLNs)**. **(A)** Representative image of FACS analyses of PLNs Foxp3+ cells in saline and *P. distasonis* gavaged mice. **(B)** Weight of spleen and pancreas of female NOD mice (n=6/group) **(C)** Number of cells in spleen and PLNs of female NOD mice **(D)** Representative image of FACS analyses of dendritic cells (CD11b-CD11c+ and CD11c+ CD11b+) in saline and *P. distasonis* gavaged mice. **(E)** Percent of CD11b+/-CD11c+/- population in splenic single cell subsets **(F)** Representative image of FACS analyses of macrophages (F4/80+cells) in spleen of saline and *P. distasonis* gavaged mice. **(G)** Percent of F4/80+ cells in CD11b+ cell subsets **(H)** Percent of F4/80+ cell subsets, where $SSC^{lo}Ly6C^{lo}$ represents residential macrophage and $SSC^{lo}Ly6C^{hi}$ represents circulatory macrophages. All samples in each panel are biologically independent. PLNs were obtained from female NOD mice oral gavaged with either *P. distasonis* (n=6) or saline (n=3). Data were expressed as means ± SEM. *, $P<0.05$, **, $P< 0.01$, ***, $P<0.001$. Statistical analysis was performed by two-tailed, unpaired Student's t-test.
